# Dynamical interactions among compositionally distinct protocell populations and its implications for evolution of early membranes

**DOI:** 10.1101/2025.02.01.636016

**Authors:** Souradeep Das, Ruchira Pal, Sudha Rajamani

**Affiliations:** Department of Biology, Indian Institute of Science Education and Research, Pune-411008, India

## Abstract

The spontaneous self-assembly of single chain amphiphiles (SCAs) would have resulted in multiple protocell species in an early-Earth niche. Considering the heterogeneity inherent in a prebiotic milieu, interactions between physicochemically distinct protocell populations was evaluated to discern if emergent properties occurred at a systems level. This study demonstrates that depending on the physicochemical properties of the membrane, interacting populations are endowed with varied emergent properties owing to their coexistence. In a multispecies paradigm involving a two-candidate protocell system, the fitter population acted as a predator and grew at the expense of the less-fit prey population. The observed growth could be attributed to the predator attaining a more robust membrane via chemical evolution. Importantly, the prey population also accrued emergent properties like molecular crowding, and coexist in balance with the predator population, without being completely outcompeted. When extrapolating these results to a three-candidate population, the outcomes were multipronged. These findings suggest a possible route for protocell membrane evolution that could have occurred even in the absence of any sophisticated protein machinery, benefitting coexisting populations, while also illustrating evolutionary trajectories that potentially resulted in functionally complex protocells.

## Introduction

Membranes are fundamental to providing a boundary condition and would have endowed early compartmentalized system with ‘selfness’,^1,2^ an indispensable criterion to undergo Darwinian evolution^3–9^. During the origins of life (OoL), protocell membranes are thought to have been mainly composed of single chain amphiphiles (SCA), such as fatty acids and their derivatives, along with other related species. These molecules comprise the prebiotically plausible amphiphilic inventory^10,11^. While the evolution of membranes to the complexity that is seen today is thought to have involved a protracted and gradual process^12^, it is pertinent to recognize that the prebiologically relevant SCA membranes could also behave like biomembranes despite their simplicity, with important implications for life’s origin and its early evolution. However, how a minimal protocell could imbibe biomimetic properties even in the absence of any sophisticated protein machinery remains an interesting problem.

SCAs show a thermodynamically-driven tendency for spontaneous self-assembly, resulting in encapsulation and concentration of solutes present in the external bulk environment. This spontaneity, in conjunction with the aforementioned compositional heterogeneity, translates to the imminent possibility that a variety of chemically distinct ‘protocellular species’ would have been readily present in the same early-Earth niche^13,14^. This would be analogically comparable to what is seen in a biodiverse ‘ecological niche’ where different kinds of organisms occupy the same environmental habitat. Inspired by the interactions that is facilitated in such ecological niches, a multispecies approach to studying physicochemically distinct protocell populations present together seemed imperative. In our experimental paradigm, we refer to such a scenario as a heterospecies population wherein the presence of protocells of different physicochemical characteristics in terms of their membrane, in a prebiotic ‘niche’ is made plausible. However, very little exploration has been undertaken to understand what systems level benefit(s) could have been possible in such interacting liposomal protocells. These systems level interactions could have either been synergistic or antagonistic, based on thermodynamic and environmental constraints acting on the protocell populations. Pertinently, this paradigm allows one to evaluate a crucial aspect, i.e., understanding whether the interaction dynamics could have had important implications for protocell ‘behaviour’ in the context of the prebiotic niche that they occupied.

In this backdrop, we aimed to characterize the population dynamics of prebiotic vesicles to discern if there was any functional advantage for the ‘fitter’ protocell population in such a heterospecies scenario. The potential benefits in such a multispecies scenario, is compared versus the well-established SCA model system that usually involves the use of a single kind of protocell, as is the case in most previous OoL studies. This paradigm of studying only a single chemical species of protocells, has been referred to as monospecies scenario in this study. The term ‘niche’ here signifies a scenario where physicochemically distinct protocellular populations are all present in a similar environment, experiencing the same pH, salt concentration, temperature etc. Therefore, in this experimental paradigm, there should be no inherent bias for a certain population in the niche to begin with, as all the environmental parameters they experience are the same. Presumably, the different protocell populations present in here are functioning at their optimum under these equilibrium conditions. Consequently, any new process/event that would occur within this experimental framework, would then solely be driven by the interacting membranes of the protocells.

In this study, we used membrane compositional heterogeneity as a metric and the resultant growth as a proxy, to show how the ‘fitter’ protocells benefit in a heterospecies scenario, while also demonstrating a mechanism for protocellular vesicle growth in such a scenario. In this paradigm, the ‘fitter’ population acts as the ‘predator’ while the ‘less fit’ population acts as the ‘prey’. Importantly, this behaviour emerges even in a simplistic two-population framework, solely due to the differences stemming from their membrane composition. Additionally, we also observed that the ‘predator’ protocell population underwent shape deformation to result in oblong and prolate vesicles, further elongating into long thread-like structures, with a continuous vesicular lumen in some cases. We were also able to discern a collateral advantage conferred on to the ‘prey’ population in our reactions. Pertinently, the prey population showed an increase in molecular crowding via a shrink-wrap mechanism^15^. This is an exciting outcome as molecular crowding has been shown to have important consequences for OoL related prebiotic reactions similar to how it is crucial for biochemical activities in an extant cell^16–19^. Studies relevant to OoL have also shown that molecular crowding increases the catalytic activity of RNA, results in differential behaviour of proteins and also affects the reaction rates of prebiotically pertinent reactions^16,20–23^. This synergistic outcome in our heterospecies scenario resulted in both the interacting populations being endowed with emergent properties while coexisting in the same niche. Therefore, we also explored a three-population system and observed that the ‘prey’ population was still acting primarily as the ‘prey’, while the other two ‘fitter’ populations were growing at its expense (**Figure 1**). The results from this study highlight the importance of understanding how synergistic processes during life’s origin would have charted the course for its subsequent evolution. It also provides crucial insights for developing a bottom-up framework for discerning a coherent approach for studying protocell ‘communities’, wherein many chemically distinct species would have contributed towards the evolution of the system in an integrated manner. In all, this study makes a strong case for thinking about it not as a singular entity but rather as a collection that behaves as a synergistic consortium of interacting cells.

**Figure 1.**
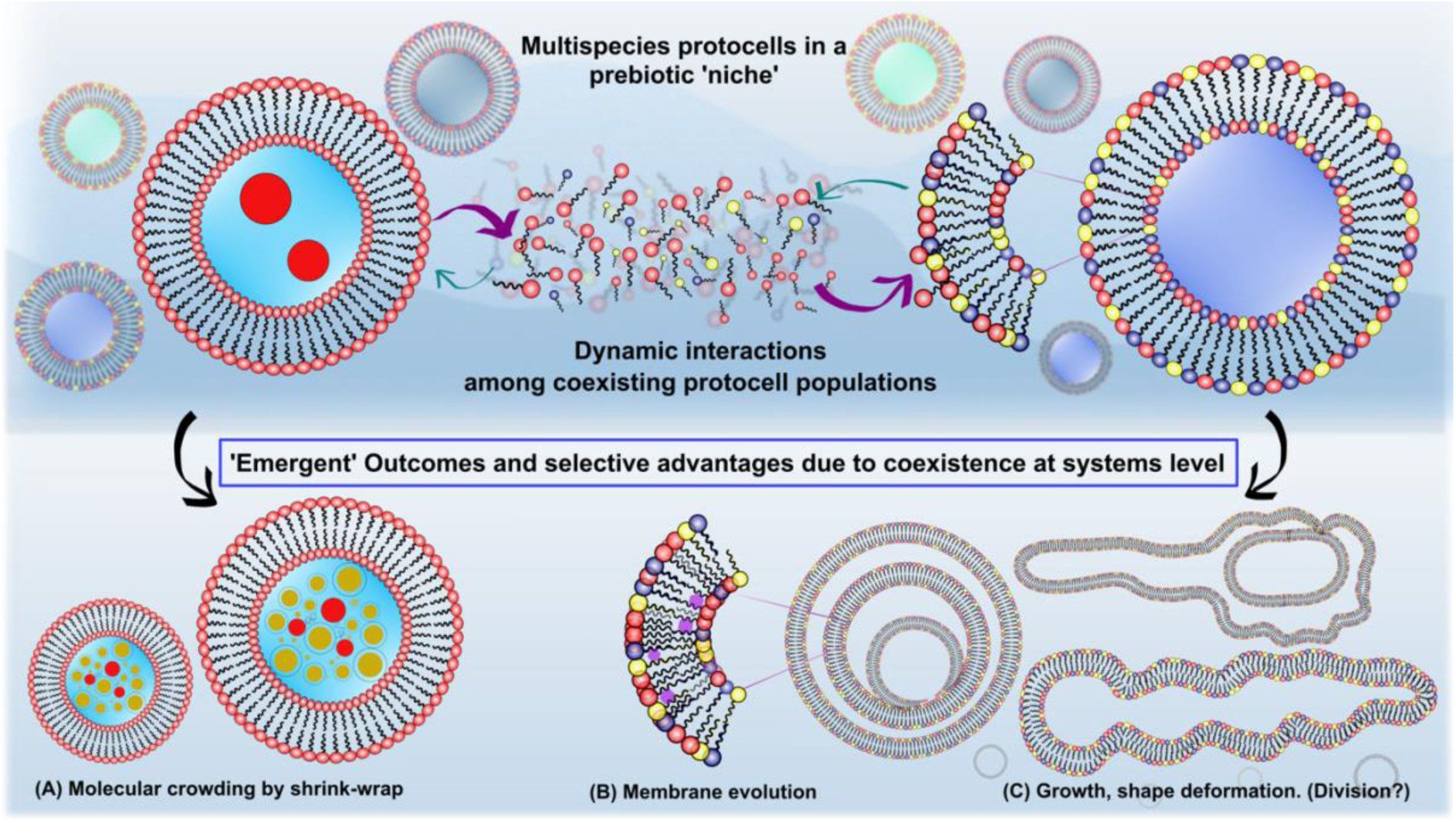
Graphical illustration of multiple protocell populations interacting dynamically in a prebiotic ‘niche’. Bottom panel shows the emergent outcomes due to such interactions driven by the distinct physicochemical properties of the membranes of the different protocell populations (heterospecies paradigm) present in the niche.

## Results

Liposomes are considered nonequilibrium supramolecular aggregates. Previous studies have shown that when the vesicle is made up of only one type of amphiphilic component, the resultant vesicles of different sizes remain distinct and kinetically trapped, without any change even over many days^24^. The model protocells in our experiment were made up of prebiotically relevant single chain amphiphiles (SCAs). Specifically, we chose a well-studied model system of C18 fatty acid and its derivative amphiphiles to represent our interacting pool of protocell populations to also situate our understanding in the context of previous related studies. The C18 amphiphiles used to generate the library of vesicles in this study were oleic acid (OA), oleyl alcohol (OOH) and glycerol monooleate (GMO) and these were the primary membrane forming moieties in our two-component and three-component reactions (**Supplementary Figure S1**). We fabricated the tertiary three-component heterogenous system using OA:OOH:GMO (abbreviated as ‘Ter’), while the two binary systems were made up of OA:OOH and OA:GMO, respectively (abbreviated as ‘Bin’). The single-component homogenous vesicle population, also the control population in our study, was comprised only of OA (abbreviated as ‘Hom’). All the aforementioned vesicle populations were prepared using dry film hydration method in 200 mM bicine buffer of pH 8.5, at room temperature. Under these conditions, all these protocellular systems are known to independently form vesicles spontaneously^25,26^, where they tend to remain as such under stable equilibrium conditions. Given this, the working hypothesis we started out with is that the SCA systems remain in equilibrium state between the free monomers in the bulk solution and the membrane embedded aggregate form. Nonetheless, upon interactions between multiple populations of distinct properties, the ‘fitter’ ones among the populations under study, would have the selective advantage as they can uptake resources more readily from the transient pool of free monomers in the bulk solution, which could result in emergent properties like memebrane growth etc (**Supplementary Figure S2**). Pertinently, upon systematic characterization, it was observed that interaction between chemically distinct protocells in the same niche, does indeed result in interesting outcomes or emergent properties. The term ‘emergent’ here defines any new property, including in structure, behaviour or function, which a vesicle system is newly endowed with.

### 2.1 System-level changes in ‘heterospecies’ vs ‘monospecies’ scenario

To detect possible changes in our reactions, we started out by testing our hypothesis by comparing the ‘heterospecies’ vs ‘monospecies’ protocell scenarios using a turbidity assay.. Turbidity changes can give preliminary insights about the nature of overall changes that could be occurring in an experimental system hosting different protocell populations mixed in a common ‘niche’. Two of the model systems were used here; a heterogenous tertiary system (Ter) and a homogenous oleic acid-based vesicle population (Hom). For setting up the turbidity assay, the two populations were prepared with two different size distributions, i.e., one large and one small population, respectively. In principle, the larger population would be the one that primarily contributing towards the measured turbidity; hence the size difference could help in identifying which population was potentially resulting in a change observed in the interacting systems.

Upon mixing with the smaller-sized homogenous OA vesicle population, the larger population comprising the heterogenous Ter population showed a gradual increase in turbidity over a course of ∼10 mins. Since turbidity is an overall readout of both the populations, to determine which population was actually causing this turbidity change, size profiles of the two populations were flipped. Notably, when the homogenous population was larger in size, it showed a decrease in turbidity (**Supplementary Figure S3**). As a consequence, the trends in turbidity values measured in these two aforementioned experiments showed a mirror pattern (**Figure 2a, Supplementary Figure 4a-c**), suggesting that one of the two protocell populations in the heterospecies scenario was responsible for the original turbidity change observed. These results indicated that the tertiary population was possibly growing while the homogenous population was diminishing in size in the heterospecies experimental scenario.

**Figure 2.**
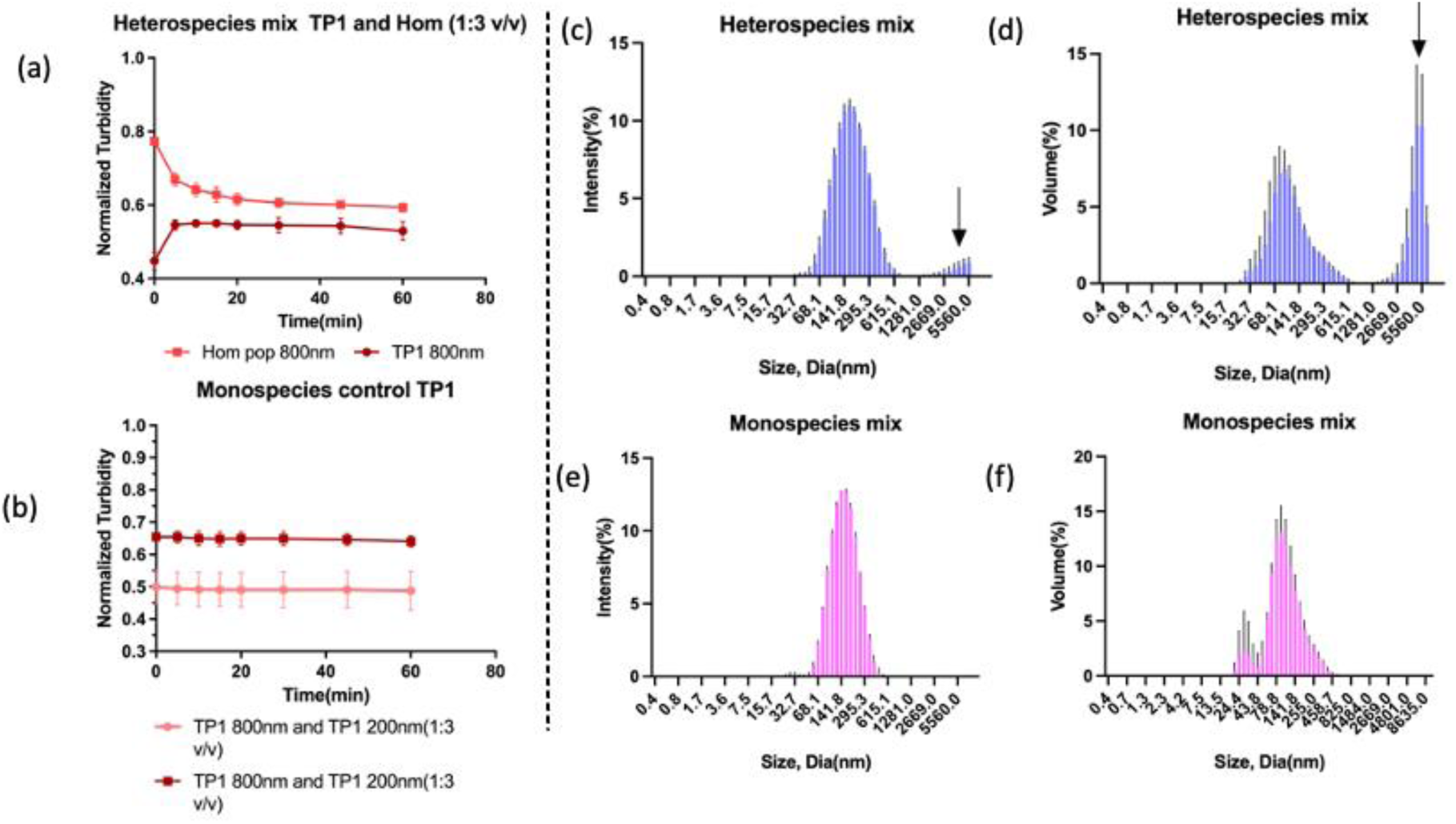
(a) Turbidity assay showing increase in turbidity over time when the larger sized Ter population (TP1) is mixed with smaller sized Hom population (Dark red trace, TP1 800nm); an exact reverse trend is observed when the Hom population is larger in size, indicating that it is becoming smaller when mixed with smaller sized Ter population (Bright red trace, Hom pop 800nm). N=3. (b) No turbidity change is observed in the systems when both the interacting populations have a similar membrane composition. This was the case even when their size difference and concentrations were varied. (c-d) Gaussian distribution of vesicles from DLS data in a heterospecies mix of ‘Ter’ and ‘Hom’ population in 1:3 v/v ratio. The diameter of the Ter vesicles was capped at 400 nm, while the size of the Hom population was kept at 50 nm. The distribution profile shows the emergence of a growth peak (indicated by arrow in c,d), which is small in intensity but large in volume. (e-f) Size (intensity) and volume distributions show no such change in a monospecies scenario. The DLS distribution plots are averaged over replicates of N=3

Various control experiments were performed using the monospecies scenario, wherein the two interacting protocell populations were physicochemically similar. In these control experiments, no obvious change in turbidity was discernable (**Figure 2b, Supplementary Figure S4d-e**). Given that all the other physical parameters between the interacting populations, were the same, the turbidity results indicated that just the simultaneous presence of chemically distinct populations was resulting in the observed turbidity change (**Figure 2a**). To confirm this, experiments were undertaken in the same paradigm albeit with different ratios (v/v) of the Ter and Hom protocell populations, while also differing the molar ratios of amphiphiles present in the Ter vesicle system. A Ter system that comprised of OA:OOH:GMO in 10:1:1 ratio also showed similar trends as was observed in the 4:1:1 case, suggesting that this phenomenon was dependent primarily on the membrane composition rather than the concentrations used in the experiment (**Supplementary Figure S4a-c**).

Dynamic light scattering (DLS) gives a robust means of quantifying system-level changes in the demographics of the particles under observation, that could be occurring in an interacting population of entities, like in the two-population heterospecies framework discussed above.. Therefore, DLS was used to investigate changes that may be occurring in the population distribution in our reactions, by looking for any upshifting of the size distribution peaks etc. To monitor the changes in our heterospecies reaction paradigm, populations of two different sizes, similar to the premise used in the turbidity assay, were prepared. The Hom and Ter populations were mixed thoroughly and incubated for 10 minutes. Subsequently, the initial and final distribution of the vesicles in the were compared, before and after mixing, to check whether any characteristic ‘growth’ peak appeared post mixing. As expected from the turbidity results, a peak with a small intensity (of ∼2-3%) showed up at around ∼4-5 μm size in the heterospecies scenario (**Figure 2c**). This growth peak was however, absent in the monospecies control and was also not discernable in the initial ‘parent’ vesicle population. Since, the size distribution of the initial population was capped well below micron size diameter by extrusion (capped at 400 nm diameter maximum) and did not undergo any mixing with any other population (**Figure 2c-f, Supplementary Figure S5a-d**).

Remarkably, the volume distribution of this peak that appeared at ∼4-5 μm, showed that these sized vesicles contributed to a large volume in the sample despite being present in very small numbers (**Figure 2d**). Overall, this indicated that a small fraction of vesicles that was present at ∼4-5 μm also had occupied a large volume when compared to several small vesicles that were present in the mixture. Appropriate controls with the monospecies scenario showed no such growth peak in the DLS size distribution (**Figure 2e-f**). Even though there would be different sized vesicles even in the monospecies scenario, with presumably varying lamellarity and/or concentrations, the fact that they had the same membrane composition failed to elicit any discernable growth peak(s) in the system. These results confirmed that growth was indeed occurring in the heterospecies scenario and it was significant enough that it changed the Gaussian distribution of the resultant vesicle population (**Figure 2c-d**). Overall, these results were very encouraging and suggested that in our heterospecies reaction paradigm, the Ter protocells were indeed being endowed with an emergent dynamic property, which was that of growth.

### 2.2 Morphology changes and growth of the ‘fitter’ population

Growth of a protocell typically means the growth of its membrane, which results in an increase in the overall volume of the resultant compartment. Incidentally, modern cells also grow in a similar manner by expanding their membranes, which occurs when they incorporate lipids synthesized within the cell^27,28^. Also, the spontaneous growth of SCA-based vesicles that occurs due to interactions with micelles, and their subsequent division, is a well-documented phenomenon^29,30^. In order to characterize protocell morphology changes, we used the same model population systems as described in the previous section to probe this and related phenomena. Two sets of experiments were performed; one to visualize and understand the changes that were occurring in the aqueous lumen of the vesicles, and another one that allowed us to evaluate the membrane component. In the first case, calcein, a water-soluble dye (**Supplementary Figure S6**), was encapsulated inside the ‘Ter’ vesicle lumen. In the latter experiment, the same Ter vesicle population was tagged with a membrane-soluble probe (either NBD-PE or LR-DHPE, as indicated). This approach allowed us to monitor the behaviour of only the heterogenous population as the size of the Hom population was diffraction limited (100 nm).

The results obtained demonstrated that the heterogenous system was indeed growing at the expense of the homogenous system. This corroborated with the trend lines that were observed in the turbidity assay as well. The vesicles grew initially to a large spherical shape and also showed visible black patches or gaps in the lumen, while keeping their shape intact (**Figure 3c-d**). These initial spherical vesicles further extended to result in long tubular continuous vesicles and this was dependent on the concentration of the available Hom population for the Ter system to ‘feed’ on (**Figure 3e-f, Supplementary Figure S7**). From these observations, we postulated that in a multispecies scenario, the heterogenous Ter population seemed to be acting as the ‘predator’ while the Hom population was behaving like the ‘prey’. Pertinently, the extent of growth observed and the shape change/deformation seen in the ‘predator’ population, seemed to depend on the availability of the ‘prey’ population. The resultant growth observed only in Ter vesicles, when mixed with the Hom vesicles, indicated that this population interaction-induced emergent phenomenon is potentially being driven by the inherent differences in the biophysical properties of the interacting model membrane systems. From a previous study in our lab, and from other relevant studies^26,31–33^, it is known that tertiary and homogenous systems have different biophysical properties in terms of their micropolarity, membrane packing, etc.^18^ Pertinently, the growth seen in our multispecies paradigm also occurred when similar concentrations of both the ‘prey’ and ‘predator’ were admixed. This further solidified our postulation that the resultant growth stemmed mainly from the populations being chemically distinct species.

**Figure 3.**
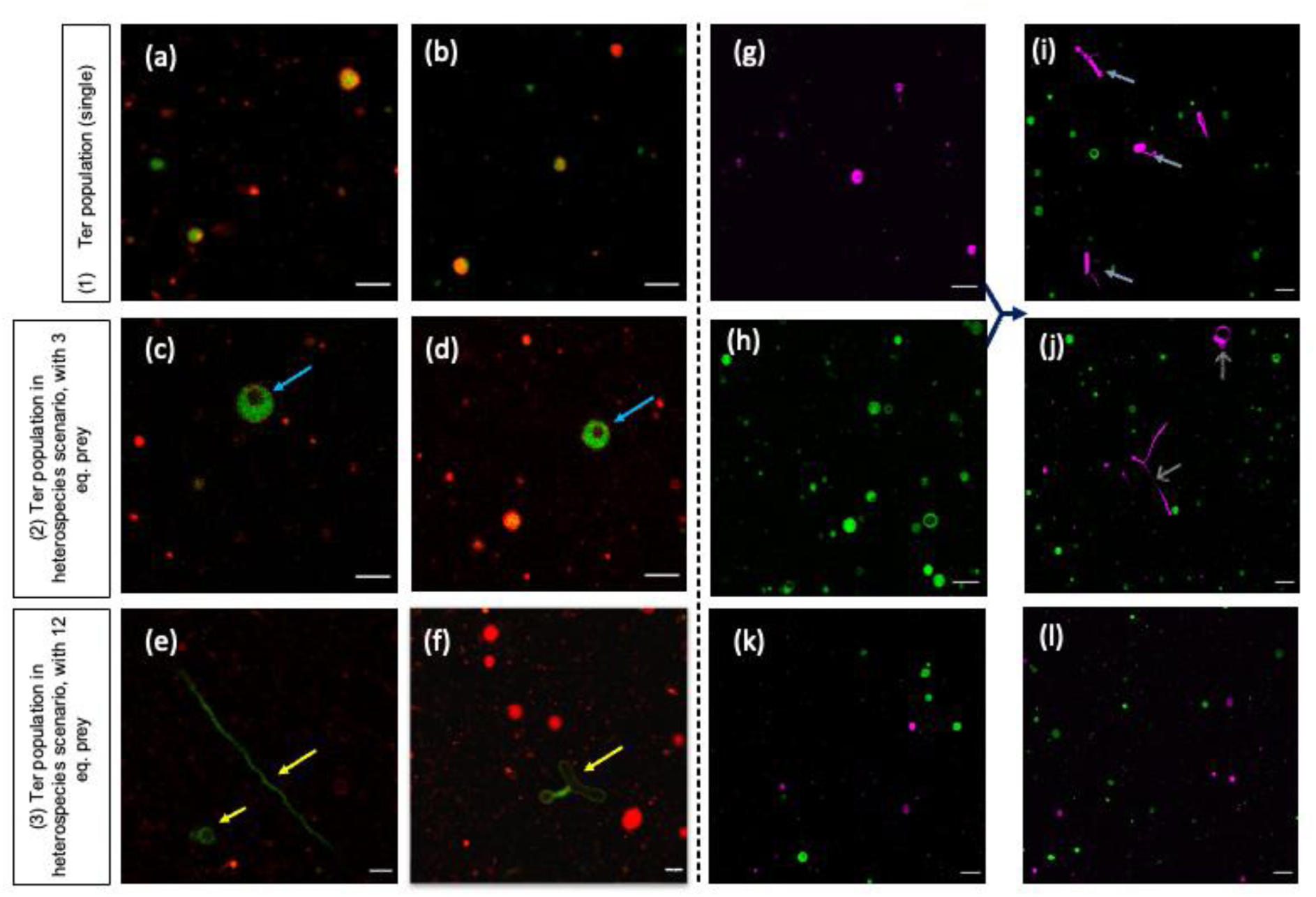
(a-b) Shows Ter parent population without any mixing. (c-d) Shows growth in Ter population due to introduction of 3eq of ‘prey’ Hom population. (e-f) Shows increased growth of Ter in presence of 12 eq of ‘prey’ population. (g-h) Image of parent vesicles from a differential tagging experiment; parent Ter ‘predator’ population and parent Hom ‘prey’ population were tagged with LR-DHPE and NBD-PE, respectively. (i-j) Shows the result of mixing the aforementioned parent populations (g and h populations) invoking a heterospecies scenario, grey arrows show growth in Ter ‘predator’ population. (k-j) Shows same experiment carried out using monospecies scenario where no growth occurred. Scale bar is 10 μm.

While the aforementioned experiments confirmed the occurrence of growth in the ‘fitter’ population, we wanted to further systematically characterize the fate of both the populations involved in this two-population experimental paradigm. To achieve this, both the interacting populations were differentially tagged, which allowed for simultaneous tracking of their respective ‘fates’. The heterogeneous population was tagged with LR-DHPE while the homogenous population was tagged with NBD-PE (**Figure 3g-h**) and both the populations were capped at same size of 5 μm by extrusion. When these two multilamellar cell-sized vesicle populations were incubated together (for 10 min at least), it was again observed that the Ter vesicle membrane grew into a range of shapes and resulted in a large distribution of sizes (**Figure 3i-j**); sometimes growing to about 12-15 times larger than the starting population size (**Figure 4a-b, Supplementary Figure S6c**). On the contrary, in the monospecies scenario, the populations being monitored remained at a similar size regime as the starting versicle pool, without any sign of changes or growth (**Figure 3k-l, Supplementary Figure S8**).

**Figure 4.**
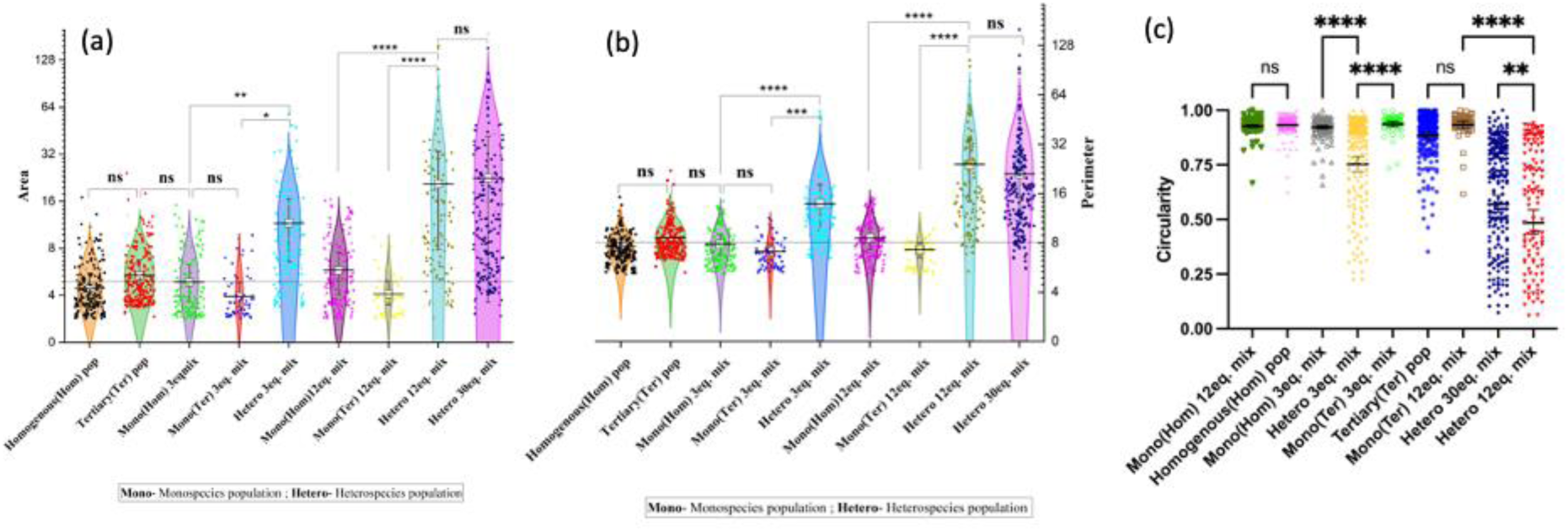
(a-b) Scatter plot of all the vesicle areas and perimeters quantified from the microscopic images. They show increase in area in a heterospecies scenario in a concentration dependent manner based on the availability of the ‘prey’ population. (c) Scatter plot of all the vesicle circularities, with 1 being a perfect circle and 0 being a complete loss of circularity. An increase in the spread indicates a higher concentration of ‘prey’ causes more loss in the normal spherical shape of the Ter vesicle, converting into more ellipsoid to long tubular structures. Each dot represents a single vesicle; for all the data sets at least 200 vesicle data was considered.

When the Ter population was incubated with different concentration equivalence of the ‘prey’ population, it was observed that growth and shape deformation readily occurred across all combinations evaluated (**Figure 4a-c**). Upon quantification of the microscopy images, the maximum change was observed when the ‘prey’ was in excess of 12 equivalence (in final concentration) than the ‘predator’ population. This corresponded to a 10-to-20-fold increase that was observed in both the area and the perimeter of the resultant Ter vesicles that showed growth in the heterospecies scenario (post mixing and incubation). Nonetheless, this phenomenon seemed to have a stalling point as well. This was clear when the prey population’s concentration was increased to 30 equivalence and there was no significant increase in growth beyond what was observed at 12 equivalence (**Figure 4a-b**). Notably, there was still an increase in the shape deformation observed in the Ter populations under this condition (Fig. 4c). Using both membrane and lumen tags, it was also confirmed that not only was the membrane growing, but rather the whole of the protocellular structure was growing without vesicle rupture due to the underlying phenomenon (**Supplementary Figure S7**).

On comparing the results obtained thus far with the detailed set of control experiments, along with the turbidity data, it was clear that no obvious growth-related changes were observed when the interacting protocell populations were compositionally the same i.e., with similar physicochemical properties. This was the case across all the vesicle combinations tested, while there was difference from 3 to 30 equivalence in the concentrations between the interacting populations in a monospecies scenario. Importantly, it did not matter even when there was an obvious size difference and probable difference in osmolarity, lamellarity, membrane tension etc. between the populations present in the monospecies scenario.. Also, when the control experiments in monospecies scenario were performed using differentially tagged vesicles (one population was tagged with LR-DHPE, another with NBD-PE), no growth or significant changes were observed as well upon microscopic analysis (**Figure 4a-b, Supplementary Figure S7**). It should be noted that there, however, was a qualitative decrease in the absolute number of vesicles present in the ‘prey’ population. This could be because of not all of the prey vesicles were potentially getting consumed by the predator population during the growth process.

### 2.3 Understanding the mechanism and causality of the observed growth phenomena

In order to understand the mechanistic underpinnings of the growth process occurring in the heterospecies scenario, we undertook systematic experiments to characterize the growth kinetics using Förster Resonance Energy Transfer (FRET) and stop-flow measurements. In previous studies, the growth kinetics of SCA vesicles when fed by micelles, was discerned using vesicle growth as a proxy^30,34^. However, we hypothesize that the mode of protocell growth in a heterospecies scenario could be different from the classical vesicle growth mechanics that occurs when vesicles are fed with chemically similar micelles^27^. This is primarily because vesicle and micelle systems are subjected to different thermodynamic constraints, thereby creating an inherent bias towards incorporation of micelles into vesicles. However, this is not the case in our reactions as all the systems are vesicles in our experimental paradigm and are present in thermodynamically ‘ideal’ conditions. Hence, to delineate the phenomenon that was at play in our interacting systems, we turned to FRET assay, which relies on a FRET pair of dyes that show a FRET-based fluorescence emission when they are in close proximity. In our heterospecies scenario, we incorporated the FRET pair (LR-DHPE and NBD-PE) in the Ter vesicles wherein the decrease in FRET efficiency (***ɛ***) suggests membrane surface growth, which is linearly related to change in the surface area due to growth.

A standard curve was first generated using known FRET dye concentrations and the corresponding ‘***ɛ***’ (see supplementary section S5, **Supplementary Fig S9**). This standard curve was later used to determine the change in the surface area (ΔSA) of the resultant vesicles and this ΔSA was plotted over time to get the rate of growth in our reaction systems. As for the main experiment involving the heterospecies scenario, both the donor and acceptor dyes were doped in the Ter system at a concentration of 0.2 mol% in total. The result obtained was a kinetic trace that showed an initial rise, which eventually plateaued, with the majority of growth happening in the first 2 minutes itself (**Figure 5a**). The kinetic plot was fitted into a single exponential curve and showed a single-phase kinetics with a rate constant of 0.033/s. This is significantly slower and distinct from the double exponential kinetics that was previously observed in the vesicle-micelle growth kinetics paradigm^3^ (**Figure 5a**). In the control experiment involving monospecies interactions, the kinetic trace obtained was a flat line, corroborating with the previous results that we obtained using turbidity and DLS, which confirmed that no growth was occurring across system, in this scenario (**Figure 5b**). It was made sure that the FRET dye by itself was not introducing any bias in the results by performing several appropriate controls (**Supplementary Figure S9**).

**Figure 5.**
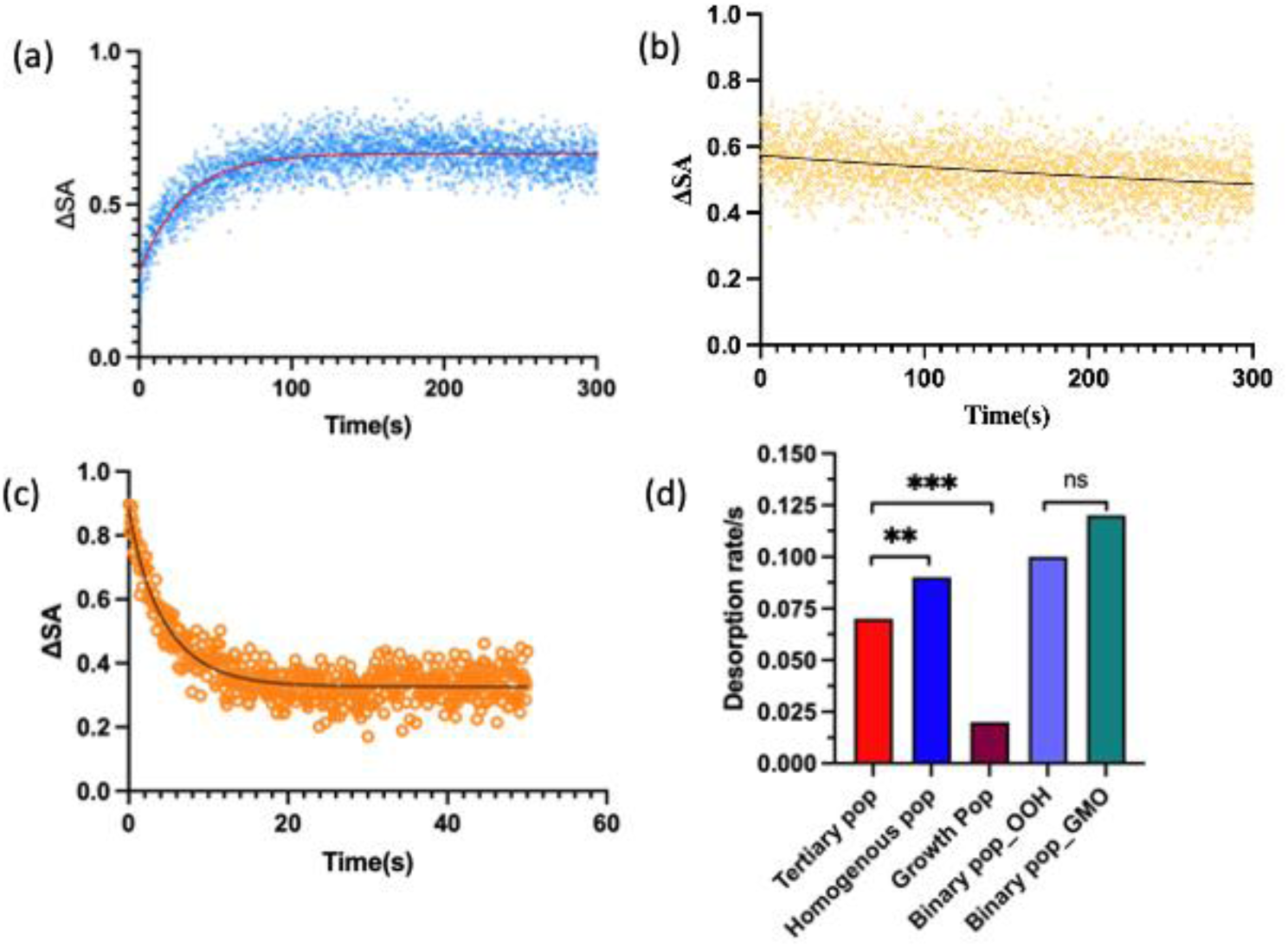
(a) FRET kinetics trace of ‘predator’ Ter population when mixed with equal amount of ‘prey’ Hom population, invoking a heterospecies scenario. ΔSA indicates the change in surface area in the Ter population (from the initial size to the final state of the vesicle). The fitted curve (red trace) depicts single exponential fit with a rate constant of 0.033/sec. (b) No significant change in ΔSA is observed in a monospecies scenario. The fitted curve (black trace) shows a linear fit. (c) FRET kinetics trace of ‘prey’ Hom population shows a decrease in ΔSA in a heterospecies scenario indicating shrinking of the Hom population. The fitted curve (black trace) shows a double exponential kinetics, depicting a fast rate constant of 0.9/sec and the slow phase with a constant of 0.2/sec. The scatter plot of the data is average of N=3 replicates. (d) Calculated desorption rates of different populations (as indicated on X axis) using HPTS assay. Growth pop (population) indicates the desorption rate of the resultant population obtained post mixing of Ter and Hom populations in a heterospecies mix. The desorption rate is calculated from the average of, N=3 replicates of HPTS assay traces (Supplementary Figure S10).

An important mechanistic reason resulting in the observed effect could potentially stem from the difference in desorption rates across the different protocell systems that were used in this study. The monomers of SCA membrane systems are known to remain in a dynamic equilibrium between the minor ‘free’ monomer form in the bulk solution and the major vesicle-incorporated form, due to the constant flip-flop of SCA monomers in and out of the membranes^34,36,37^. In a scenario where multiple populations are present, the aforesaid phenomenon would benefit those protocell populations which have an intrinsically lower monomer escape rate, thereby resulting in a higher retention of the SCA within those membranes. We performed an assay to test this SCA monomer escape rate (or desorption rate) using an HPTS encapsulated POPC reporter vesicle. Using the stopped-flow setup in the fluorimeter, the test SCA population was mixed at pH∼8.5 with the POPC reporter vesicle that had an internal lumen pH of 7.1. Being a pH sensitive probe, the change in the fluorescence of HPTS would directly correlate with the monomer escaping from the SCA vesicle system and entering the POPC reporter. This results in proton efflux out of the POPC reporter vesicle due to the flip-flop movement of the SCAs that have a dissociable head group.

From the above experiment it was found that the monomer escape rate or desorption rate in the Hom population was the highest at 0.09/sec. In comparison, the ‘predator’ Ter population showed a desorption rate of 0.065/sec. The rates observed for both the binary systems of OA:OOH and OA:GMO, was around 0.1/sec (**Figure 5d**). These results provide preliminary insights into why the thermodynamics of the system is favoring the ‘fitter’ Ter population in the heterospecies scenario. We also tried to test the desorption rate of the resultant vesicle population that was obtained after growth had resulted in this paradigm. This was done by separating the resultant population from the starting population, by using a size-based filtration protocol (see supplementary section S6). Notably, the population that resulted from growth in the heterospecies scenario (noted as growth population), did show the lowest desorption rate of 0.02/sec. This is more than 3 times lesser than the parent predator population of Ter vesicles and 4.5 times lesser than the Hom prey population. This finding indicated that the population that resulted post-growth, showed the maximum monomer retention. This was potentially by attaining a composition post growth that actually changed its membrane property significantly (**Figure 5d, Supplementary Figure S10**), thereby stalling the process at this putative ‘ideal’ composition’ that it could attain under the study conditions described. As we had postulated in our initial hypothesis, this desorption rate-driven kinetics was possibly one of the major driver of the growth that we observe in our heterospecies experimental paradigm.

### 2.4 Change in the biophysical properties of membranes resulting from growth

Given the FRET kinetics and desorption rates, fast dynamics of the SCA membrane system seemed to play a decisive role in the phenomena under study here. Although membranes are thought to be dynamic in general, SCA membranes are known to be many folds more dynamic than conventional phospholipid membranes^37,38^. This is true when considering the movement of the monomers; either within the bilayer or their flip-flop in between the bilayer and also their escape into the bulk solution. This stems from the transient pseudo-diacyl structures that SCAs form around their pK_a,_ unlike what is seen in their true diacyl counterparts^5,37^. Therefore, interactions of SCA membranes with their surrounding environment is known to affect the protocells’ fitness in multiple ways^26,39^. This dynamicity also allows them to spontaneously respond and modulate themselves to environmental stochasticity, making them an ideal protocell model system for rapid chemical evolution. A primitive mode of growth and proliferation has been demonstrated in a cell wall-less organism called L-form bacteria, which is very similar to what has been previously shown using certain model protocells in the OoL field^28,30^. Such growth is typically achieved by an increase in the surface-to-volume ratio resulting from increased membrane synthesis, with division being subsequently triggered by random blebbing due to Rayleigh instability. An important driver of this phenomenon is membrane order, which can be readily measured using generalized polarization (GP) of the membrane.

In the aforementioned backdrop, we wanted to characterize the any change in physicochemical property of the membranes due to growth in our study using a solvatochromic probe called Laurdan that gives us information about the GP value of the membranes. The GP kinetics correspond to the ordered or disordered phases in a membrane and how this shift over time. A negative GP value of around -0.6 was obtained for the model OA-based SCA vesicles^18^, suggesting that the membrane was disordered with higher water accessibility due to less packing of the bilayer. The growth phenomena observed in the heterospecies scenario resulted in an exponential increase in the GP value of the predator tertiary population (**Figure 6a**). This meant that the Ter population membranes were becoming more ordered, accounting for the observed growth, even when there was no inherent bias in the system to begin with. Although the increase in GP value showed a trend that was similar to the growth kinetics, the rate of change in GP value had slower temporal kinetics. Interestingly, we also observed that the GP value of the prey population was increasing in a heterospecies experimental scenario (**Supplementary Figure S12a**). This is possibly being driven by the escape of monomers from the membrane, which could also exclude water from the bilayer, thereby increasing the GP of the prey population.

**Figure 6.**
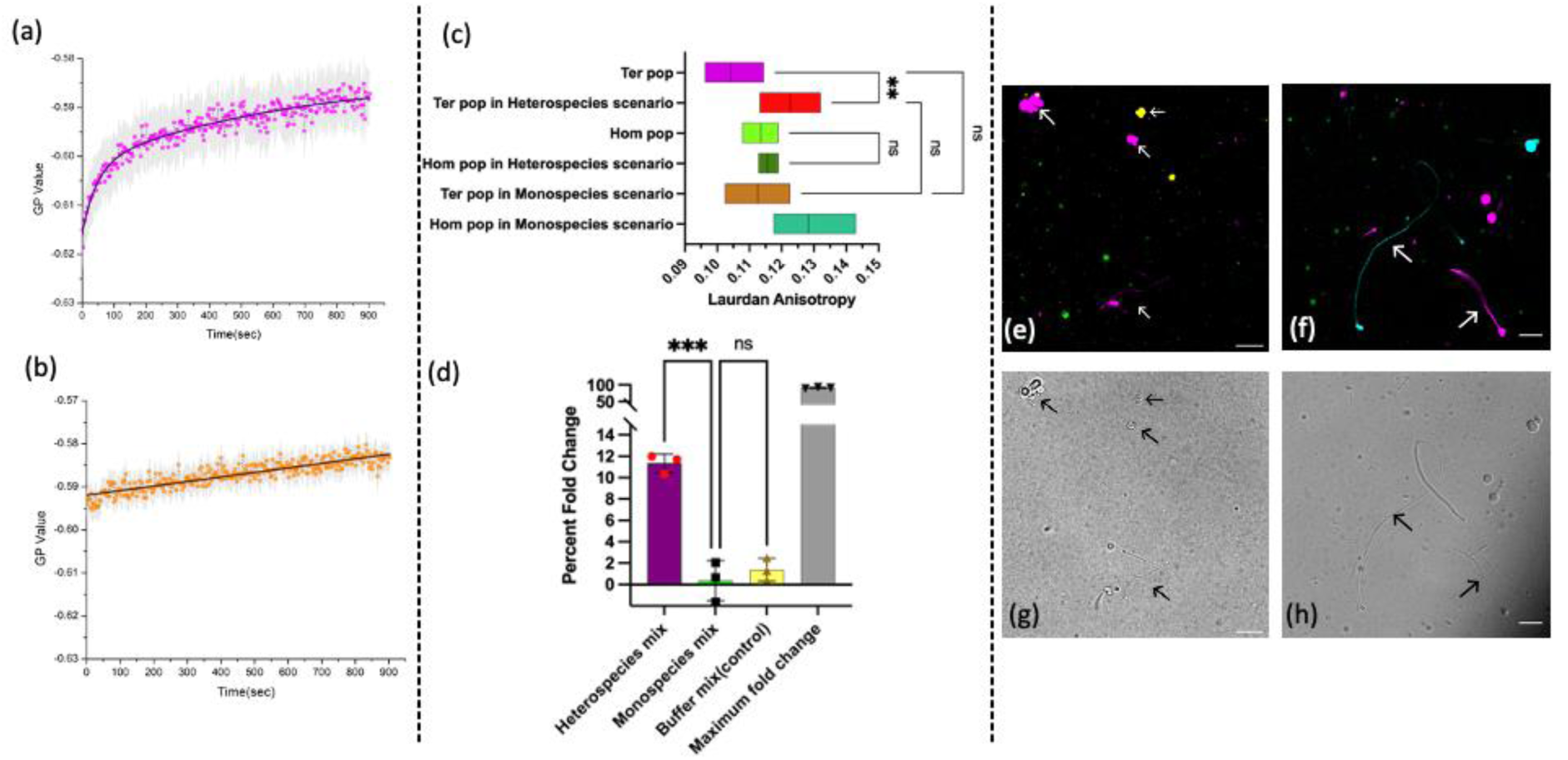
(a-b) GP kinetics trace of ‘predator’ Ter population in a (a) heterospecies scenario vs a (b) monospecies scenario, showing that GP shifts towards positive value in Ter population happens only in heterospecies scenario owing to increase in membrane order due to growth. N=3. (c) Membrane anisotropy changes in Ter and Hom pop is plotted in Heterospecies vs monospecies scenario. N=3. (d) Calcein quenching assay showing molecular crowding occurring in the Hom population. Percent fold change indicates change in fluorescence of calcein due to concentration dependent quenching by molecular crowding between the initial and final state of the vesicle populations, N=3. (e) Showing simultaneous growth of both Ter (Magenta) and Binary 1 (Bin 1) (Yellow) populations and (f) Showing simultaneous growth of both Ter (Magenta) and Binary 2 (Bin 2) (Cyan) populations in the presence of Hom ‘prey’ population (Green). (g-h) Differential interference contrast (DIC) image of the same field as the respective top frame, showing the grown and shape deformed vesicle. Scale bars are 10μm.

It is known that phospholipid-based membranes are kinetically trapped and not nearly as dynamic as SCA membranes, with a GP value that is much more positive^18^. Hence, we ran a positive control using POPC vesicles to discern how the GP would shift in this case. When mixed with a SCA Hom vesicle population, POPC vesicles absorb monomers from the SCA vesicle population resulting in an expected decrease in GP value (**Supplementary Figure S12e**), which is a known phenomenon^35^. In case of control experiments involving the interaction of two SCA-based monospecies populations, there was no apparent change in the GP value, which was consistent with the results from previous experiments (**Figure 6b, Supplementary Figure S12b**). These results validated the GP assay observations along with another control experiment that showed no significant changes occurring just by mixing vesicles with buffer, as a baseline (**Supplementary Figure S12c-d**). Using a similar strategy, changes in membrane anisotropy were also measured using Laurdan dye as it is an indicator of membrane order/packing, and the results obtained corroborated the observations from the GP assay data. Typically, the initial anisotropy value of a Hom population is generally more than the Ter population. However, in the heterospecies experimental scenario, the predator Ter population showed a significant increase in membrane anisotropy upon growth (**Figure 6c**). Expectedly, in the control monospecies scenario, no significant change in the anisotropy value was observed (**Figure 6c**). This further suggested that in the growing Ter population, membrane stabilization and chemical evolution of membrane was being aided due to water exclusion upon growth, which increased the membrane packing in the predator population in our heterospecies experimental paradigm.

### 2.5 Fate of the ‘prey’ population

While it became apparent from the results obtained thus far that the Ter population was growing, it was imperative to also systematically chacaracterize the fate of the Hom population in the heterospecies scenario. In previous studies, researchers have shown that usually one population was completely ‘consumed’ by the other^30,40^. However, in our studies, preliminary observations were indicative of a ‘balance in coexistence’. Notably, microscopy results showed that both the prey Hom population and the predator Ter populations coexisted, despite the predator growing at the expense of the prey and this is the case, as the observed Ter vesicle growth was not occurring indefinitely (**Figure 4a-b**). Nonetheless, both microscopy and DLS analysis were not amenable to helping in discerning what was happening to the prey population; was it really diminishing in absolute numbers or was some other event taking place? Therefore, we monitored the fate of the ‘prey’ population using a similar FRET-based assay by incorporating the previously discussed FRET pair, this time in the Hom population, while keeping all the other parameters same as in the aforementioned set of FRET experiments.

Upon rapid mixing using stopped-flow, the Hom prey population showed ‘shrinkage’ of vesicles due to a decrease in the membrane surface area (**Figure 5c**). This definitively confirmed there are two distinct dynamics in the behaviour of the different kinds of protocells that was being observed under our experimental paradigm. Curiously, the shrinkage rate appeared as a two-phase exponential curve; the faster phase had a rate constant of 0.9/sec and the slower phase had a rate constant of 0.2/sec. This is probably happening due to some of the Hom ‘prey’ population getting completely consumed after shrinking by the ‘predator’ population. The latter slow rate suggests a probable existence of a population that gradually loses the monomer and shrinks, but is still not fully consumed over the course of the reaction. Pertinently, in conjunction with the HPTS desorption assay, this result proves that the process of growth of the Ter population was indeed happening due to monomer escape from the prey population, reconfirming our initial hypothesis. The kinetics of shrinking is similar to the kinetics of growth, albeit reversed, which also further confirmed that the growth and shrinkage that were being observed in the two populations that coexist together, was indeed occurring simultaneously.

This shrinking observed in the Hom vesicles, leads to an interesting unintended consequence; that the prey population might be experiencing molecular crowding in their lumen. This kind of ‘shrink-wrapping’ has been observed previously, which is a phenomenon where a vesicle shrinks and could concentrate entrapped molecules in the lumen due to extraction of its building block monomers^15^. This study showed that fatty acid vesicles underwent ‘shrink-wrapping’ when mixed with a POPC-fatty acid (FA) mixed micelle, in which the vesicles could concentrate the entrapped molecules in the lumen. In a prebiotic context, molecular crowding has been argued to be important for facilitating relevant prebiotic reactions and for protocell evolution^20,41^. It has also been shown to increase RNA activity and to facilitate competition among protocell populations^41,42^.

Given the above, we wanted to characterize the possibility of an emergent phenomenon of molecular crowding that might arise in the Hom prey population. To study this, calcein was encapsulated in Hom population, as a proxy molecule in a concentration of 10 mM that shows partial quenching of calcein (see supplementary section, volume shrinkage assay). Inside the vesicular protocell lumen further concentration increase of calcein would allow quantifying the amount of crowding that was resulting in the Hom vesicle population. Upon addition of the predator Ter population to the prey Hom population, calcein signal showed a marked decrease in the signal due to concentration dependent self-quenching. This is a direct readout of molecular crowding occurring inside the lumen and this was being triggered only in the heterospecies scenario. The percentage fold change from initial to final calcein signal obtained has been plotted (**Figure 6d**). Pertinently, no significant difference in fluorescence was observed when the protocells were made to interact in a monospecies paradigm and this was similar to what was also seen when the populations were just mixed with equiomolar buffer solutions in negative controls that were performed to rule other unexpected contribution from calcein leakage from vesicles or osmotic shock. Upon quantification, the volume of prey population was observed to shrink by up to 6-7 folds when compared to the original volume of the starting vesicles (**Figure 6d**). This means that the Hom population is indeed shrinking and experiencing molecular crowding in the lumen when in coexistence with the predator Ter population; a happy consequence for thermodynamically challenged prebiotically relevant reactions that typically are affected by dilution issues.

### 2.6 Live imaging of vesicle growth and introducing additional complexity to the heterospecies experimental paradigm

All the microscopy analysis discussed thus far involved the use of imaging techniques that give a static overview of events that occur in our experiments. However, investigating the growth phenomenon using live imaging, should reveal a more detailed understanding of the underlying process and also hint if there was possible division occurring in the system. Since our main goal was to understand population level behavior of different interacting protocellular entities, we also wanted to further ‘complexify’ the reaction system as well. Towards this, we undertook an experiment in which there were three interacting populations in the same environmental niche. In order to monitor changes occurring simultaneously in the three different populations, live imaging was undertaken using surface tethered vesicles as it offers a robust means of deciphering potential growth and related phenomena. We initially standardized this assay using the two-population system by tethering the Ter predator vesicle population on to a glass surface (40 mm coverslip). Thereafter, the Hom prey population at 4 equivalence higher concentration was flowed using a flow-cell set up on the microscope. Images were then acquired live for a period of at least 10 minutes.

The live time lapse imaging of the process clearly demonstrated the event of growth and the associated shape change that resulted from it. Pertinently, it also showed that after a certain extent of growth had occurred, the process stalled and this was also observed from the quantification of microscopy data (see section 2.2). The resultant structures were stable over the time duration of the experiment i.e., for up to 1 hr. (Supplementary video 1, Supplementary Fig. S14). Nonetheless, no significantly noticeable division event was observed even after flowing in the ‘prey’ population for over 30 minutes. This could be potentially due to the resultant growth not happening to a point of making the structure unstable, since the amount of incoming monomers that the Ter membranes can accept seems to have a limit (section 2.2, **Figure 4a, b**). Pertinently, despite having an excess of prey population, not all of the prey Hom population was being consumed by the predator (**Figure 3c-f**), resulting in the coexistence of multiple populations under our experimental conditions. Even though within the time duration of our experiment we see no obvious membrane division, we hypothesize that given enough time and resources, the resultant long tubular structures could eventually reach an unstable state, which in previous work has been shown to be energetically unfavourable due to their inherent instability^30,40^.

In order to complexify our starting population mix, a third protocell population was introduced to the heterospecies experimental paradigm that was also differentially tagged (from the other two populations). Towards this, from the library of the four populations used in this study, one of the two binary populations (OA:OOH (Bin1) or OA:GMO (Bin2)) was added to the population mixture that already contained the Ter and Hom vesicular systems. They were simultaneously imaged using confocal microscopy as they were all tagged with different fluorophores allowing for tracking of their individual fates. On monitoring the three populations simultaneously under similar parameters of buffer, pH, temp. etc, (see methods), it was observed in case of both the binary populations that both Ter and Bin populations were growing at the expense of the homogenous population, in a 3-systems heterospecies scenario (**Figure 6e-h**). This assay demonstrated that the binary and the tertiary populations were both growing simultaneously, and seemingly, to a similar extent. Nevertheless, the morphology of the resultant structures that resulted in Bin1 and Bin2 systems were different in this three population heterospecies scenario, and this was observable qualitatively using confocal microscopy (**Figure 6e-f**).

Perceivably, the concurrent presence of the Bin populations probably affected the extent of growth that was possible in the Ter population when present in the same niche due to competition for the prey Hom vesicles. This observation further underscored that in niches with multiple vesicle populations, growth occurred in the ‘fitter’ populations, even when competing for the same set of resources. This was also corroborated by the HPTS desorption assay where the desorption rate of the Ter population was significantly lower than that of Hom population (**Figure 6c**). In all, these results highlight the occurrence of complex interactions between the three interacting populations when present in the same niche. This further alludes to the potential consequences that can result when the system is complexified even further. However, the systematic exploration of this would be experimentally non-trivial and will require modelling approaches to discern all the possible outcomes and the mechanistic underpinnings involved in even more complex scenarios.

## Discussion

The scenario of multispecies protocell population has been predicted computationally to varied extents using different lipid-aggregate based models before^32,43,44^. However, there is very little to no experimental evidence of how such an event might have happened. Overall, the emergent outcomes observed in our studies, including membrane growth, vesicular shape change and molecular crowding, highlights the importance of empirically discerning the nature of multispecies interactions in a heterogenous prebiotic niche. Once a ‘fitter’ population emerges in the ‘niche’, the equilibrium in the dynamics can shift again, in principle, allowing for the protocell membrane to gradually move towards a more functional and robust system. In this study, we set out to understand the implications of a prebiotically realistic scenario where different protocell populations would have coexisted and interacted in the same niche. We achieved this by using membrane-related processes as a proxy, wherein SCA-based model vesicle systems were allowed to interact with each other either in a monospecies or a heterospecies paradigm. It is evident from our results that the ramifications could be varied, with the outcome depending on the composition and nature of the protocell systems involved. A recent paper reported another case study on protocell population level interplay, albeit using two homogenous vesicle populations of different acyl chain lengths. The thermodynamic equilibrium of the systems experimented with in this study was different to begin with, which inherently biases one system over the other in terms of vesicle formation and sustenance. Nonetheless, it underscored the importance of discerning systematically how protocellular population interactions could have impinged on membrane evolution^40^. In our study, we characterized this aspect and demonstrated that even in the absence of any environmental or systemic bias, interactions of physicochemically distinct protocells could indeed result in an outcome that clearly illustrated an emergent phenomenon advantageous to a subset of the population involved. The possibility of multiple coexisting populations, with each showing different emergent properties, demonstrates a novel take on how the protocell population can act synergistically as a ‘whole’ system.

The introduction of a second physicochemically distinct population seemingly shifted the thermodynamic energy minima of the ‘fitter’ system. Notably, this led to changes in its membrane property, resulting in the ‘molecular evolution’ of this membrane towards a more robust membrane system in terms of its physicochemical properties when compared to the initial population. Our study also shows a finite limit up to which a membrane can uptake molecule to grow under our experimental conditions in our two-population paradigm. Despite the prey losing its monomers and shrinking, it nevertheless led to a beneficial outcome of molecular crowding being facilitated in its lumen, while it also continued to coexist in the system without getting completely diminished. On the other hand, the predator population showed an emergent property of growth and also exhibited significant shape deformation with a loss of circularity. This growth and shape change observed in the ‘predator’ population was correlated to the availability of the ‘prey’ population.

Taken together, in a two-population system, both of them benefitted from coexisting in a heterospecies scenario, an outcome that did not occur in a monospecies experimental paradigm. Pertinently, the observation of a shrinking prey population, along with the single exponential growth curve obtained for the predator population, rules out the potential formation of any intermediate aggregate as has been reported previously in an experimental paradigm involving vesicle-micellar growth kinetics. Further, a potential direct fusion event between the two interacting heterospecies populations can be ruled out, since such an event would have been slower than when growth occurs only by monomer incorporation^38,42^. If this was indeed happening in our heterospecies paradigm, such an event would have resulted in a double exponential curve, which was not observed in our reactions.

Unlike in previous studies, this growth phenomena could be solely triggered by difference in membrane composition in our studies, and even when there was no concentration difference between the prey and predator populations in the interacting niche. Nonetheless, the resultant growth became more prominent when the prey was present at a higher concentration than the predator. It is relevant to note that the loss of circularity of a vesicle (when growth results), is an energetically unfavorable process. However, since the Ter membranes were gaining significant biophysical advantages in terms of membrane properties such as, increase in membrane packing, it seemed to be able to overcome this barrier to undergo shape change in our heterospecies scenario. This eventually resulted in a ‘fitter’ membrane composition than in the initial population. And, when resources are not limiting, generation of new daughter protocells by spontaneous division of long tube-like vesicular structures, could potentially result in the generation of new vesicles given environmental variations in the surrounding bulk environment, for e.g., like pH and temperature.

While understanding the physicochemical changes that are causing the growth phenomenon using desorption assay, an intriguing feature was observed; the whole population in a heterospecies scenario became less dynamic than the initial population after achieving certain growth. This is because it was getting better at retaining the SCA monomers in the membrane, resulting in lesser escape rate of the monomers into bulk solution. Overall, it was evident from the GP assay that the interacting populations were getting more stable when compared to the parent populations, due to the membrane becoming more ordered, a property that makes membranes amenable for further evolution (**Figure 6a**). The incorporation of OA Hom monomers into the predator Ter population is potentially resulting in in better packing of the membrane, which drives the GP value in the positive direction. Significantly, the increasing tendency of the lipids to remain in the vesicles indicates that a protracted period of such an evolutionary drive can modify membrane properties towards a compositionally superior membrane system, making it more ‘kinetically trapped’.

Due to experimental constraints, it is impossible to demonstrate all possible outcomes when considering the possibility that several kinds of prebiotically plausible amphiphiles could have been present in a given early Earth niche. Nevertheless, as a first step in this direction, we incrementally complexified our system to experiment with a two and three-population scenario. The latter experiments clearly show that both the binary and tertiary populations can act as predator in a heterospecies paradigm, growing at the expense of the prey Hom population. In all, most of our preliminary observations from a two-population heterospecies scenario also held true for the three-population systems. However, further characterization of the dynamics of three-population systems, and other combinations of increasing complexity involving more chemically distinct protocell types, will be pertinent for a systematic understanding of which molecular evolutionary pathways could have resulted in highly functionally evolved cellular entities. We believe, this study can be used as a groundwork for further computational modelling of much more complex systems with more interacting populations to predict interesting outcomes and phenomena.

In conclusion, this study’s finding underscores the importance of factoring in the unlimited possibilities and permutations that would have been plausible in a molecularly heterogenous prebiotic niche, wherein ‘fitter’ membranes could have come about as a consequence of interactions between physicochemically distinct protocell populations. Observations gleaned from such studies also has implications for synthetic biology approaches involving minimal cells inspired by extant biology. Specifically, understanding the biochemical underpinnings of such interacting systems will be crucial to delineating how synergistic protocell populations could have contributed to a self-sustaining, networked population system. Studying interactions in multiple coexisting populations over protracted timescales (using modelling), could better illustrate the probable evolutionary trajectories that potentially resulted in functionally complex protocells, which could have eventually set the stage for the emergence of cellular entities that more closely resemble extant life.

## Experimental section

### Vesicle-based protocell population preparation

The SCA amphiphilies were mixed in desired ratios; for tertiary (Ter) heterogeneous population, OA:OOH:GMO were mixed in 4:1:1 ratio for most of the experiments (mentioned as Tertiary population or Ter and Tertiary population-1/TP1 specifically in turbidity assay data). For one set of turbidity assay OA:OOH:GMO were mixed in a 10:1:1 ratio (mentioned as Tertiary population-2/TP2 only in turbidity assay data). The two binary systems (Bin1 and Bin2) were prepared by mixing either OA:OOH or OA:GMO in 2:1 ratio. This kept the overall ratio between the major membrane forming component i.e. OA with all its derivatives at 2:1 at all times and the ratios were calculated as fractions of the total lipid concentration of the system (mentioned in the experiments). In most of the experiments, the lipid concentrations of the vesicle systems was kept at 5 mM. Whenever a different concentration was used, it has been mentioned accordingly in the manuscript/methods section. This lipid film was further resuspended using bicine buffer (200 mM) of pH 8.5 to prepare all the samples (unless mentioned otherwise). For further details refer to Supplementary method section S1. Details of the materials used for these and below mentioned experiments can be found in the supplementary information.

### Turbidity assay

For turbidity experiments, one protocellular vesicle population was extruded with 800 nm membrane using the Avanti mini-extruder and the other was extruded with 200 nm polycarbonate membrane. This resulted in two population sizes, i.e., one large (800 nm) and one small population (200 nm), respectively. The wavelength of the incident light used was 400 nm and the turbidity values were obtained using a Shimadzu UV-Spectrophometer UV-1800, which tells about the scattering of the particles present in the sample. In principle, the larger population comprised of the 800 nm (extruded) vesicles, should primarily contribute towards the measured turbidity since the incident light is of 400 nm wavelength (Supplementary Fig. S3). The two populations were incubated together in a quartz cuvette and readings were taken at different time points over a maximum period of 1 hour, while continuing to keep the sample in the same cuvette without introducing any disturbance. The turbidity values were normalised and plotted (Y axis) against time (in minutes, X axis) to get a qualitative idea of the changes occurring in any of our reactions.

### Dynamic Light Scattering (DLS)

DLS was performed using a Malvern Panalytical Zetasizer Nano ZS90. For DLS experiments, the heterogeneous population was extruded with a 400 nm polycarbonate membrane and the homogenous population was treated in a similar manner but was extruded using a 50 nm membrane.. Extrusion was done for at least 13 passes using the Avanti polar mini extruder. We used 1:3 (v/v) mixing of a larger population (400 nm) with a smaller population (50 nm) in experiments of heterospeies or monospecies scenario. The heterospecies populations experiments and monospecies population controls were incubated for at least 15 minutes and then data was recorded. The data was then plotted as a Gaussian distribution wherein the total area under the curve for each plot amounts to 100%. Each bar represents the amount/ percentage of the respective size vesicles that were detected in a specific size range or of a specific hydrodynamic radii (diameter in nanometres), as mentioned on the X axis. See supplementary section S3 for more details regarding this experiment.

### FRET assay

For the Förster resonance energy transfer (FRET) assay, vesicles were prepared by dry film hydration with the total lipid concentration kept at 3 mM. In addition to the SCA lipids, 0.02 mol% of the total lipid concentration was FRET dyes (NBD-PE and LR-DHPE) that were also added while preparing the lipid films for the vesicle population. First, a standard curve was obtained using known concentrations of FRET dyes. The samples were prepared separately for each data points and the FRET efficiency (ε) was measured across the different concentrations. A total of seven points for the ‘ε’ (each with three replicates) were plotted against the known concentration of FRET dyes to get the standard curve (Supplementary Fig. S9a, see section S5). All the vesicle samples for this assay were extruded using a 100 nm poly carbonate membrane to make small unilamellar vesicles (SUV). The kinetics measurements were acquired using a Horiba Fluoromax^+^, with a SFA-20 rapid mix stopped-flow accessory. For the kinetics experiment, the population under study was doped with a total of 0.2mol% of the FRET fluorophore pair while the other population to be mixed was not tagged (until otherwise mentioned). Two populations that were to be mixed were loaded on to two different sample syringes in the stopped-flow accessory. During injection, 125 µL of both ‘predator’ and ‘prey’ populations were mixed rapidly using the pneumatic-drive trigger in the attached sample cuvette to make a heterospecies or monospecies mix and the florescence were recorded using excitation and emission wavelength of 463 nm and 530 nm, respectively. The florescence intensity was then converted into change in the surface area (ΔSA) using the standard curve and an appropriate fit was generated using Graphpad Prism. Data was recorded for up to 5 minutes and plotted as needed.

### Lipid desorption assay-

A vesicle made with POPC (1-palmitoyl-2-oleoyl-phosphocholine) phospholipid was used as the reporter vesicle, which reported the fluorescence changes correlated with the lipid desorption. 3 mM POPC vesicles were prepared by dry film preparation method as before and resuspended in 20 mM pH ∼7.1 bicine buffer, along with 2 mM HPTS (8-Hydroxypyrene-1,3,6-trisulfonic acid trisodium salt) dye. The total lipid concentrations for all SCA vesicle systems were kept at 5 mM. The buffer used for the SCA vesicles was bicine buffer (20 mM), set at pH ∼8.5 and 50 mM sodium chloride (NaCl). Both the POPC reporter vesicles and the SCA protocell vesicle samples were extruded using a 100 nm filter and performed SEC (see supplementary section S2, S6) with it. It is pertinent to note that the outside pH of the POPC reporter vesicle was adjusted to pH∼8.5, to match the bulk pH of the other sample solution (SCA vesicle solution) with a base pulse using 1M NaOH solution just before using it in the experiment. The SCA vesicles were typically mixed with the reporter POPC vesicles using the stopped-flow spectrofluorometer unit (see supplementary section S6). During injection, POPC reporter and the respective SCA vesicle samples were injected (1:1 v/v) into the sample cuvette using the pneumatic drive stopped-flow accessory. The pH-sensitive HPTS emission increase was measured at 510 nm (post excitation at 460 nm). In our experiments, the HPTS showed an increase in fluorescence due to basification of the lumen by incorporation of SCA molecules into POPC reporter vesicles.

### Vesicle shrinkage assay

10 mM calcein, was encapsulated in the Hom population made of OA lipids of 20 mM concentration and extruded with a 200 nm membrane to characterize how much crowding as happening in these vesicles (see supplementary section S7). The ‘predator’ Ter population was prepared using a total lipid concentration of 20 mM, using the dry film hydration method. After performing SEC, the eluted vesicle fractions were pooled and just before the experiment, 100 μM of cobalt chloride (CoCl_2_) was added to the vesicle sample. Calcein encapsulated Hom population was added to the ‘prey’ Hom populations and the reaction was monitored via calcein fluorescence using an excitation/emission combination of 460/515 nm, respectively, with the same stopped-flow setup as mentioned before. The change that resulted due to crowding that was facilitated in the Hom population’s lumen, occurred within 10 seconds after using the stopped flow device. The fold change in fluorescence between the initial and final fluorescence readout was plotted after the reaction reached equilibrium. After equilibrium, at least 10 data points were averaged to make sure that we captured the completion point of the reaction.

### Laurdan GP and anisotropy assay

The sample preparation for the GP kinetics assay was done using the dry film hydration (see supplementary section S7). along with addition of Laurdan dye as the fluorescent probe of choice because of its solvatochromic nature. The laurdan dye was incorporated into only those samples whose membrane dynamics was under study. Excitation/ emission wavelength was 350 nm and emission wavelengths were recorded at 440 nm and 497 nm. Reaction mixes of Hom and Ter vesicle populations was monitored for 15 minutes (900 seconds) upon mixing. GP values were calculated using the formula *GP = (I _440_ – I _497_)/ (I _440_ + I _497_),* where *I* is the intensity values at the respective wavelengths (see supplementary section S8), which correspond to the two intensity peaks of Laurdan in ordered vs disordered phases. The change in GP value in a system under consideration has been plotted against time. We also fitted curves for all the experiments (**Supplementary Table 1**).

Laurdan anisotropy single-point measurement was also recorded for Ter and Hom populations in a heterospecies scenario. This was done as a steady state measurement comparing ‘before and after’ change in anisotropy value (see supplementary section S9).

### Preparation of surface tethered vesicle-

The protocol used was modified and adapted from two previously published articles using the 40-mm Bioptech coverslip^45,46^. For passivation, these coverslips were incubated in molten PEG (95% PEG 4000 and 5% 5K-PEG-Biotin w/w, Sigma-Aldrich) at 90℃ for 48 hrs. For this experiment the vesicle populations were made as described previously in ‘vesicle-based protocell population preparation’ section (see supplementary section S1). Along with the lipid, 18:1 Biotinyl Cap PE was added in 0.02 mol% of the total lipid concentration to the Ter population, while making the dry lipid film. The Ter population was also tagged with octadecyl rhodamine B (R18, at 0.02 mol%) for effective visualization, and the total concentration of lipid was kept at 5 mM. The concentration of Hom population was kept at 20 mM. The intended vesicle was tethered onto the passivated coverslip by biotin – streptavidin-biotin interaction. The experiments were performed using a FCS2 closed chamber flow-cell attached with a microperfusion low-flow pump (Bioptechs) under an Olympus IX83 inverted microscope (see supplementary section S10 for more details). The Hom prey population was flowed into the flow cell at a final concentration of 20 mM and live images were taken over a period of 10 minutes using a LED light source with an excitation wavelength of 550 nm (for red channel). Images were taken at an interval of 1 second to visualize changes in the tethered populations while flowing in the ‘prey’ Hom population.

## Acknowledgements

SR acknowledges research support from IISER Pune, Department of Science and Technology’s Science and Engineering Research Board (SERB) (Grant No. CRG/2021/001851) and Department of Biotechnology, Govt. of India (Grant No. BT/PR43193/BRB/10/2006/2021). SD would like to thank CSIR, India for fellowship support. SD would also like to thank Dr. Soumya Bhattacharya and Prof. Thomas J. Pucadyil for help with live imaging experiments. The authors would like to thank Prof. Manickam Jayakannan for access to DLS instrument. We are thankful to IISER Pune microscopy facility for access to the microscopes and technical help. We also like to thank COoL lab members for their critical comments and inputs.

## Author Contributions

S.D. and S.R. conceived the project and designed the study. S.D. designed the assays, performed most of the experiments and analyzed the data. R.P. carried out the Laurdan GP shift assay and anisotropy experiments. S.D. made all the illustrations. S.D. and S.R. wrote the manuscript.

